## Supplementary Figures for "Neural correlates and determinants of approach-avoidance conflict in the prelimbic prefrontal cortex"

### SUPPLEMENTARY FIGURE 1

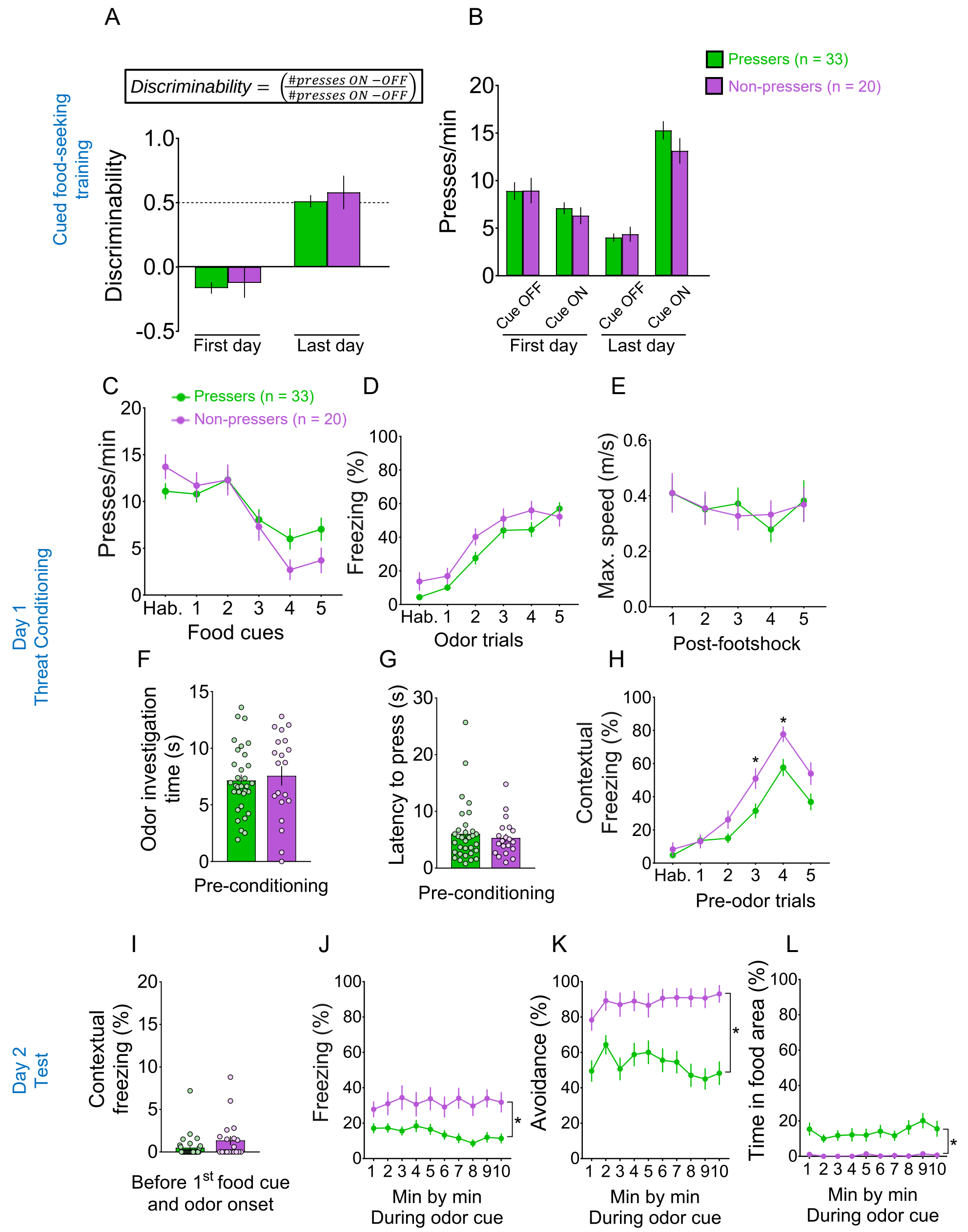

**Supplementary Figure 1. *Pressers* and *Non-Pressers* showed similar behavioral responses during cued food-seeking training and olfactory threat conditioning. (A)** *Pressers* and *Non-pressers* learned to discriminate equally the food cue off from the food cue on periods during the training phase (Group, $F_{(1, 55)} = 2.166$; $p = 0.146$, Time, $F_{(1, 55)} = 238.7$, $p < 0.001$, Interaction $F_{(1, 55)} = 0.106$, $p = 0.745$, Bonferroni *post-hoc* for early vs. late $p < 0.001$). Discriminability was calculated as the number of lever , pressers during the cue on period (30 s) minus the number of lever presses during the cue off period (30 s before cue) divided by the total number of presses during both periods. **(B)** *Pressers* and *Non-Pressers* showed the same lever press rates during the cue off and cue on periods across the cued food-seeking training ($F_{(3, 153)} = 1.28$, $p = 0.283$). **(C-E)** *Pressers* and *Non-Pressers* exhibited the same (C) lever press rates ($F_{(1, 51)} = 0.265$, $p = 0.608$), (D) freezing levels ($F_{(1, 51)} = 3.737$, $p = 0.058$), and (E) maximum speed ($F_{(1, 51)} = 6.538e007$, $p = 0.999$) during the olfactory threat conditioning training. **(F-G)** *Pressers* and *Non-Pressers* spent the same (F) time investigating the odor (Unpaired Student's t-test, $t = 0.43$, $p = 0.665$) and showed the same (G) latency to press the lever (Unpaired Student's t-test, $t = 0.55$, $p = 0.578$) before the first odor-shock pairing. **(H)** Non-Pressers showed higher contextual freezing during the third and fourth pre-odor trials (30 s before odor onset), compared to *Pressers* ($F_{(5,250)} = 3.038$, $p = 0.011$; Bonferroni *post-hoc* test, third pre-odor trial: $p = 0.0214$; fourth pre-odor trial: $p = 0.015$). **(I-L)** During the test session on Day 2, *Pressers* and *Non-Pressers* showed the same levels of (I) contextual freezing before the first food cue and odor onset (Shapiro-Wilk normality test, $p < 0.001$, Mann Whitney, $U = 248$, $p = 0.113$). The behavioral responses were different between groups but constant across the session for (J) freezing (Group $F_{(1,50)} = 13.07$, $p < 0.001$; Interaction, $F_{(9,450)} = 1.327$, $p = 0.220$), (K) avoidance (Group $F_{(1,50)} = 20.31$, $p < 0.001$; Interaction, $F_{(9,450)} = 2.109$, $p = 0.027$, Bonferroni *post-hoc* min 1 vs. min 10 $p > 0.999$) and (L) time in food area (Group $F_{(1,50)} = 117.5$, $p = 0.001$; Interaction, $F_{(9, 450)} = 0.573$, $p = 0.819$). Combined data for all *Pressers* and *Non-Pressers* used across the experiments. Data shown as mean ± SEM. Two-way ANOVA repeated measures followed by Bonferroni *post-hoc* test, * $p < 0.05$ for group comparison. All statistical analysis details are presented in table S1.

### SUPPLEMENTARY FIGURE 2

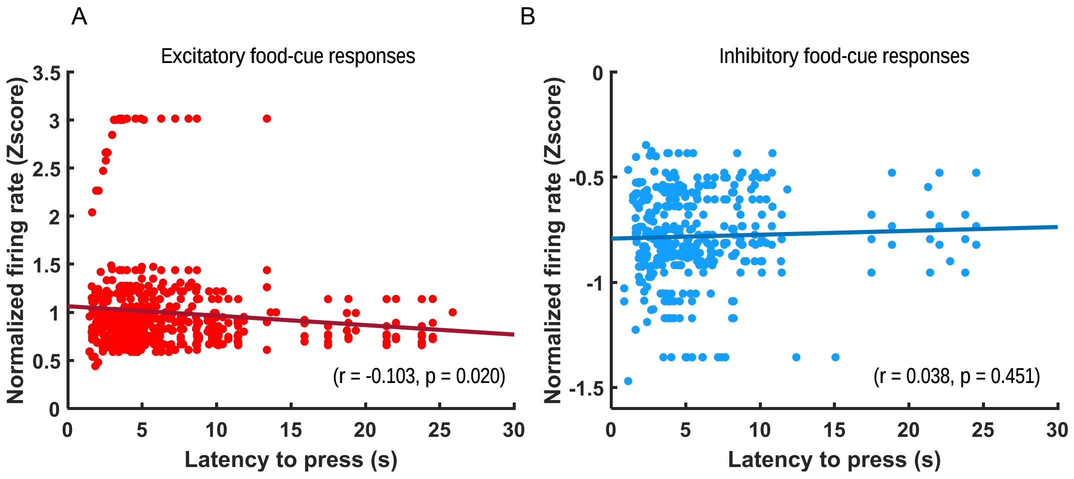

**Supplementary Figure 2. Correlation between food cue-evoked PL activity and lever press latency during the conflict phase in *Pressers*. (A)** Scatter plot showing a significant inverse correlation between the normalized firing rate (Z-score) of excitatory food-cue responsive neurons and the latency to press the lever after the food cue onset during the conflict phase ($r = -0.103$, $p = 0.020$). **(B)** Scatter plot showing lack of correlation between the normalized firing rate (Z-score) of inhibitory food-cue responsive neurons and the latency to press the lever after the food cue onset during the conflict phase ($r = 0.038$, $p = 0.451$). Each data point denotes the averaged Z-scored response of one neuron until the animal pressed the lever (Y axis) vs. the respective latency to press during each food-cue presentation (X axis). The black lines represent the linear regression of the data points, fitting the first degree polynomial to the data (see Methods for details). The equation for the linear fitting in the excitatory case is $y = 0.0098*x + 1.1$ and for inhibitory cases is $y = 0.0018*x - 0.79$.

### SUPPLEMENTARY FIGURE 3

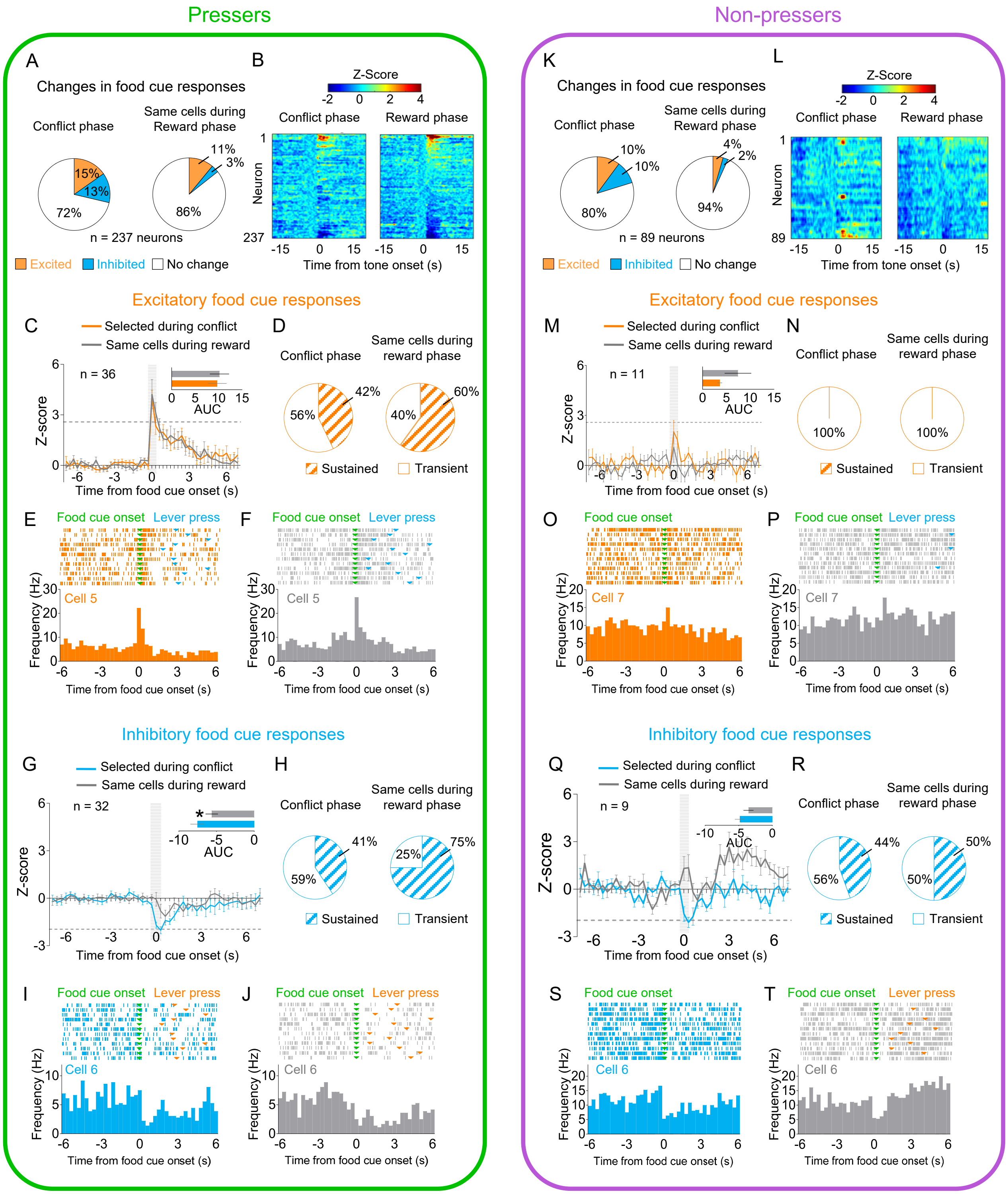

**Supplementary Figure 3. Changes in PL responses to reward cues selected during the conflict phase for *Pressers* and *Non-Pressers*. (A)** Pie charts showing changes in PL firing rate in response to food cues during conflict (left) vs. reward (right) phases for *Pressers* (n = 237 neurons from 25 rats, Fisher Exact Test, responsive during conflict phase: n = 68 , responsive during reward phase: n = 34, $p < 0.001$, excitatory during conflict phase: n = 36, excitatory during reward phase: n = 26, $p = 0.22$; inhibitory during conflict phase: n = 32, inhibitory during reward phase: n = 8, $p < 0.001$). **(B)** Heatmap of Z-scored neural activities for PL neurons selected during conflict phase and tracked to reward phase. **(C)** Average peri-stimulus time histograms (PSTHs) for all PL neurons showing excitatory food cue responses (Z-score > 2.58, dotted line) during conflict (orange line) compared to the same cells during reward (gray line). C, Inset: Differences in the positive area under the curve (AUC) between the two phases (Wilcoxon test, W = 122, excitatory responses reward phase vs. conflict phase, $p = 0.346$). **(D)** Pie charts showing the percentage of sustained vs. transient excitatory food-cue responses in PL neurons during the conflict phase with the same neurons tracked back during the reward phase. **(E-F)** Representative PSTHs for a PL neuron showing excitatory responses to food cues during the (E) conflict phase vs. the same neuron during the (F) reward phase. **(G)** Average PSTHs for all PL neurons showing inhibitory food cue responses (Z-score < -1.96, dotted line) during conflict (blue line) compared to the same cells during reward (gray line). G, Inset: Differences in the negative AUC between the two phases (Wilcoxon test, W = 266, inhibitory responses reward phase vs. conflict phase, $p = 0.011$). **(H)** Pie charts showing the percentage of sustained vs. transient inhibitory food-cue responses in PL neurons during the conflict phase with the same neurons tracked back during the reward phase. **(I-J)** Representative PSTHs for a PL neuron showing inhibitory responses to food cues during the (I) conflict phase vs. the same neuron during the (J) reward phase. **(K)** Pie charts showing changes in PL firing rate in response to food cues during conflict (left) vs. reward (right) phases for *Non-Presses* (n = 89 neurons from 7 rats, Fisher Exact Test, responsive during conflict phase: n = 18, responsive during reward phase: n = 6, $p = 0.0129$, excitatory during conflict phase: n = 9, excitatory during reward phase: n = 4, $p = 0.2486$; inhibitory during conflict phase: n = 9, inhibitory during reward phase: n = 2, $p = 0.0573$). **(L-M)** Same as B-C, but for *Non-Pressers*. M, Inset: Differences in the positive AUC between the two phases (Wilcoxon test, W = 19, excitatory responses reward phase vs. conflict phase, $p = 0.300$). **(N)** Pie charts showing the percentage of sustained vs. transient excitatory food-cue responses in PL neurons during the conflict phase with the same neurons tracked back during the reward phase. **(O-Q)** Same as E-G, but for *Non-Pressers*. Q, Inset: Differences in the negative AUC between the two phases (Wilcoxon test, W = 11, excitatory responses reward phase vs. conflict phase, $p = 0.570$). **(R)** Pie charts showing the percentage of sustained vs. transient inhibitory food-cue responses in PL neurons during the conflict phase with the same neurons tracked back during the reward phase. **(S-T)** Same as I-J, but for *Non-Pressers*. For all Shapiro-Wilk normality test, $p < 0.05$. All statistical analysis details are presented in table S1.

### SUPPLEMENTARY FIGURE 4

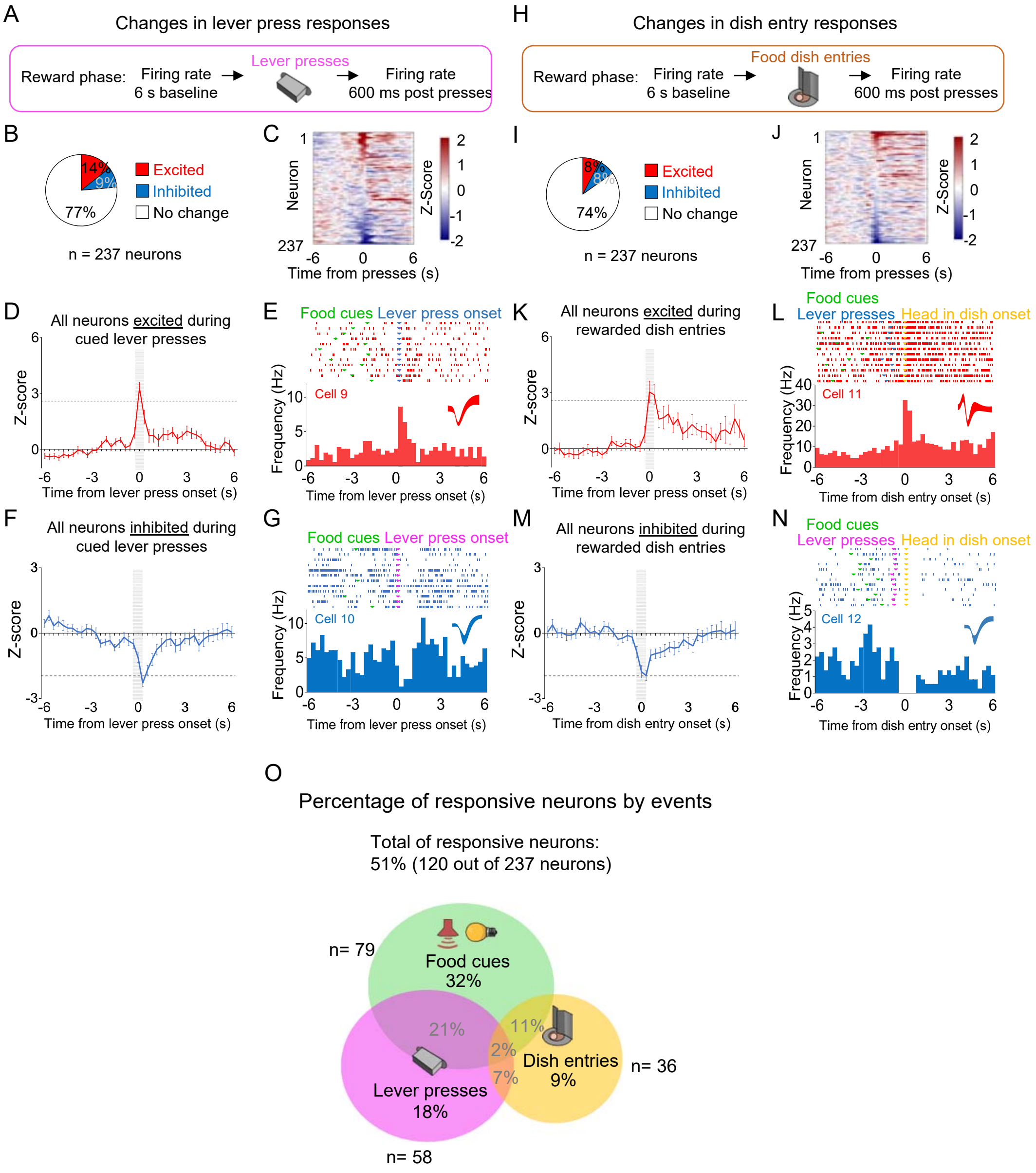

**Supplementary Figure 4. Distinct subpopulations of PL neurons change their firing rates in response to food cues, lever pressers, and dish entries.**

**(A)** Schematic of the recordings during cued lever presses. **(B)** Pie charts showing the percentage of lever press responsive neurons during the reward phase. **(C)** Heatmap showing the normalized firing rate of individual PL neurons time-locked for lever presses. **(D)** Average peri-stimulus time histogram (PSTH) of all PL neurons showing excitatory lever-press responses. **(E)** Raster plot and PSTH of a representative PL neuron showing excitatory lever-press responses. **(F)** Average PSTH of all PL neurons showing inhibitory lever-press responses. **(G)** Raster plot and PSTH of a representative PL neuron showing inhibitory lever-press responses. **(H)** Schematic of the recordings during rewarded food dish entries. **(I)** Pie charts showing the percentage of dish entry responsive neurons during the reward phase (n = 237 neurons). **(J)** Heatmap showing the normalized firing rate of individual PL neurons time-locked for rewarded food dish entries. **(K)** Average PSTH of all PL neurons showing excitatory dish entry responses. **(L)** Raster plot and PSTH of a representative PL neuron showing excitatory dish entry responses. **(M)** Average PSTH of all PL neurons showing inhibitory dish entry responses. **(N)** Raster plot and PSTH of a representative PL neuron showing inhibitory dish entry responses. **(O)** Venn Diagram showing the percentage of all responsive neurons (120 out of 237) by events. Most of the responsive neurons (59%) responded exclusively to one of the events. n = 237 neurons from 25 rats.

### SUPPLEMENTARY FIGURE 5

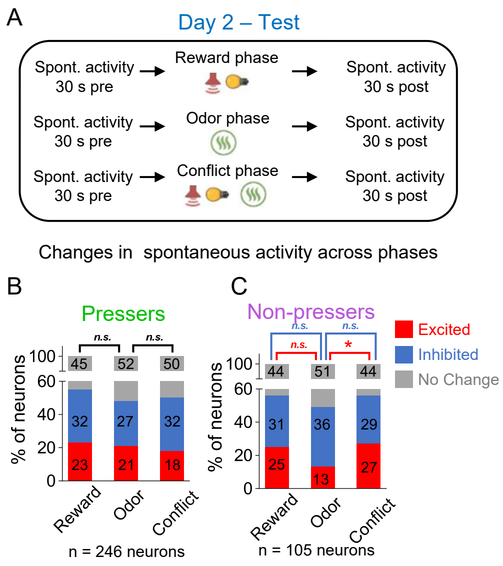

**Supplementary Figure 5. Changes in PL spontaneous activity in *Pressers* and *Non-Pressers* across the different phases of the test session. (A)** Timeline of PL recordings for spontaneous activity during test. **(B-C)** Stacked bars showing the percentage of PL neurons that changed their spontaneous firing rates across the different phases of the test in both *Pressers* (B) and *Non-Pressers* (C). *Non-Pressers* showed a significant increase in the proportion of excited neurons from the odor to the conflict phase (Fisher Exact Test, excited from Reward phase to Odor phase: 14 neurons, excited from Odor phase to Conflict phase: 28 neurons, $p = 0.015$. n.s. = non-significant, * $p < 0.05$. All statistical analysis details are presented in table S1.

### SUPPLEMENTARY FIGURE 6

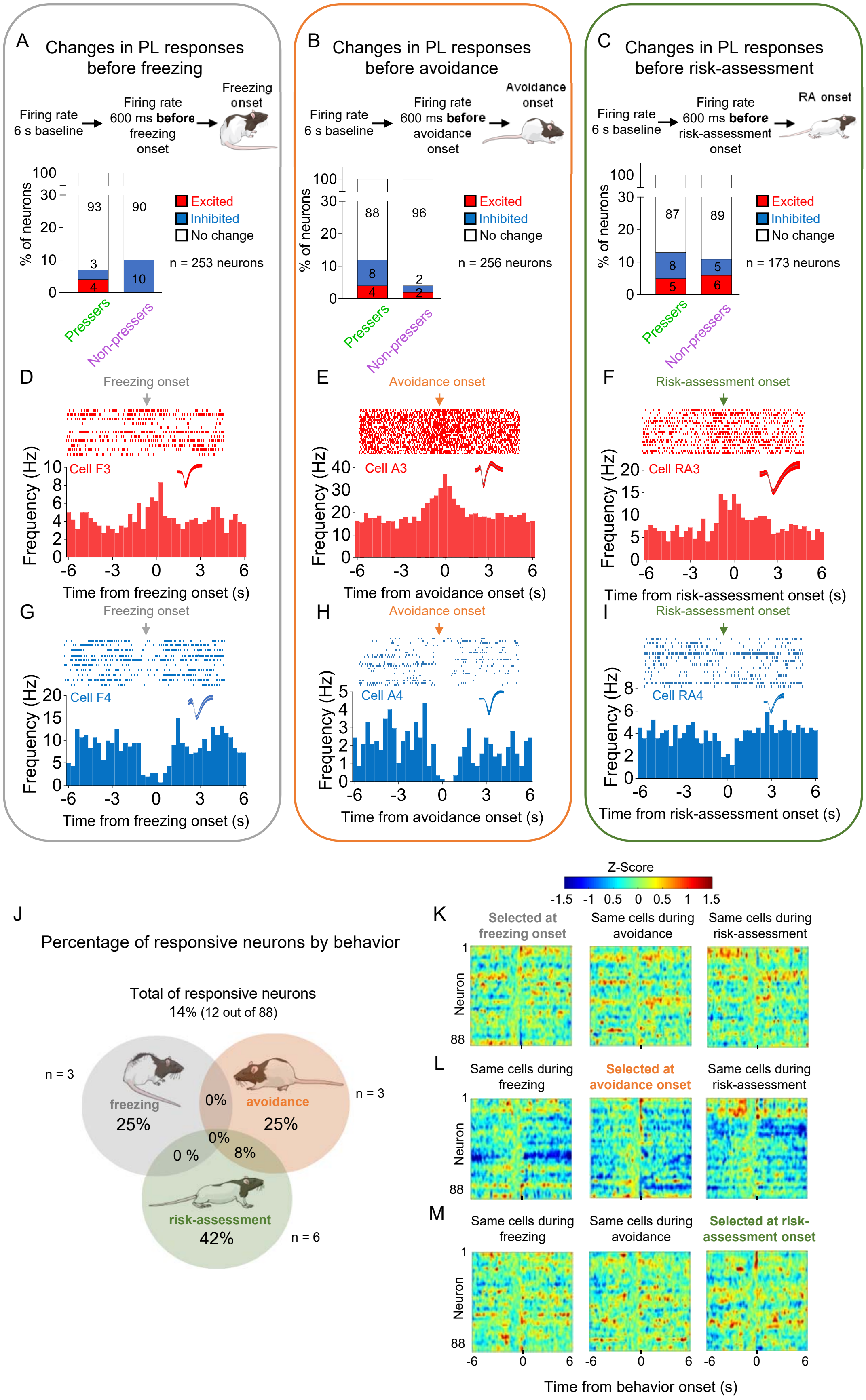

**Supplementary Figure 6. PL activity anticipates the onset of freezing, avoidance or risk-assessment behaviors in both *Pressers* and *Non-Pressers*. (A-C)** Both *Pressers* and *Non-Pressers* showed the same number and proportion of excitatory and inhibitory PL responses 600 ms before the onset of (A) freezing (Fisher Exact Test, $p = 0.34$), (B) avoidance (Fisher Exact Test, $p = 0.1249$) or (C) risk-assessment (RA, Fisher Exact Test, $p = 0.8168$) behaviors. **(D-F)** Representative PSTHs for distinct PL neurons showing excitatory responses 600 ms before the onset of (D) freezing, (E) avoidance or (F) risk-assessment behaviors. **(G-I)** Representative PSTHs for distinct PL neurons showing inhibitory responses 600 ms before the onset of freezing (G), avoidance (H) or risk-assessment (I) behaviors. **(J)** Venn Diagram showing the percentage of all PL responsive neurons (12 out of 88 neurons) by behavior. Most of the responsive neurons responded selectively at the onset of one of the behaviors. **(K-M)** Heatmap of Z-scored neural activities for PL neurons selected at 600 ms before the onset of freezing (K), avoidance (L) or risk-assessment behavior (M) with the same cells tracked during the other behaviors. The threshold used to identify significant differences per neurons was Z-score > 2.58 for excitation and Z-score < -1.96 for inhibition. Stack bar values were compared using Fisher Exact Test, n.s. = non-significant. All statistical analysis details are presented in table S1.

### SUPPLEMENTARY FIGURE 7

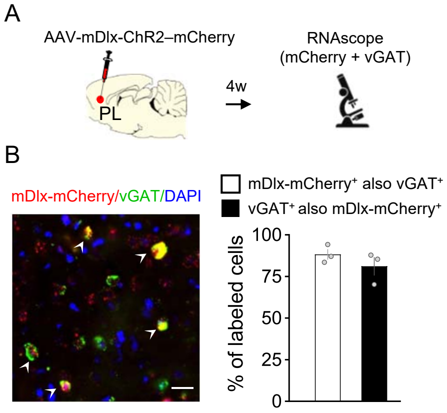

**Supplementary Figure 7. Validation of the mDlx promoter used for viral vector targeting of GABAergic neurons in PL. (A)** Schematic of AAV-mDlx-ChR2–mCherry infusion and fluorescent *in situ* hybridization in PL. **(B, left)** Representative micrograph of PL cells expressing mRNA for mCherry (red label), the GABAergic marker vGAT (green label), or both (white arrowheads). Blue label: DAPI. **(B, right)** Quantification of the total of cells expressing mDlx-mCherry that were also labeled with vGAT (white bar), or cells labeled with vGAT that also expressed mDlx-mCherry (black bar). n = 3 rats.

### SUPPLEMENTARY FIGURE 8

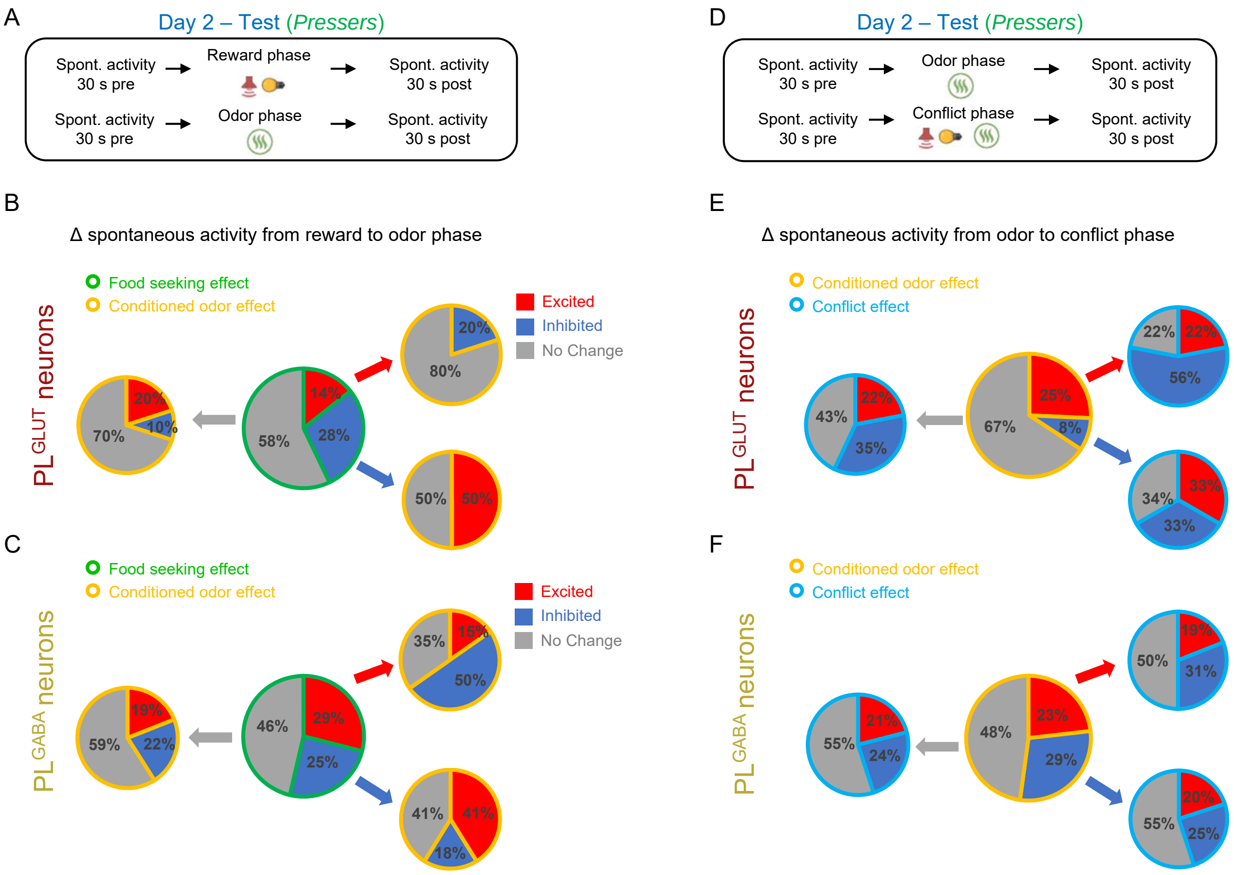

**Supplementary Figure 8. Changes in the spontaneous firing rate of $PL^{GLUT}$ and $PL^{GABA}$ neurons across the different phases of the test session. (A)** Timeline of PL recordings for changes in spontaneous activity from reward to odor phase. **(B-C)** Pie charts showing the proportion of (B) $PL^{GLUT}$ neurons (n = 36) and (C) $PL^{GABA}$ neurons (n = 69) that changed their spontaneous firing rates from baseline to reward phase (central pie charts, green borders) and from reward phase to odor phase (peripheral pie charts, yellow borders) in *Pressers*. All $PL^{GLUT}$ neurons that were excited or inhibited during the reward phase responded in opposite directions or did not respond during the odor phase. **(D)** Timeline of PL recordings for changes in spontaneous activity from odor to conflict phase. **(E-F)** Pie charts showing the proportion of (E) $PL^{GLUT}$ neurons and (F) $PL^{GABA}$ neurons that changed their spontaneous firing rates from reward to odor phase (central pie charts, yellow borders) and from odor phase to conflict phase (peripheral pie charts, blue borders) in *Pressers*. For all the pie charts the activity of the same cells was tracked across the session.

### SUPPLEMENTARY FIGURE 9

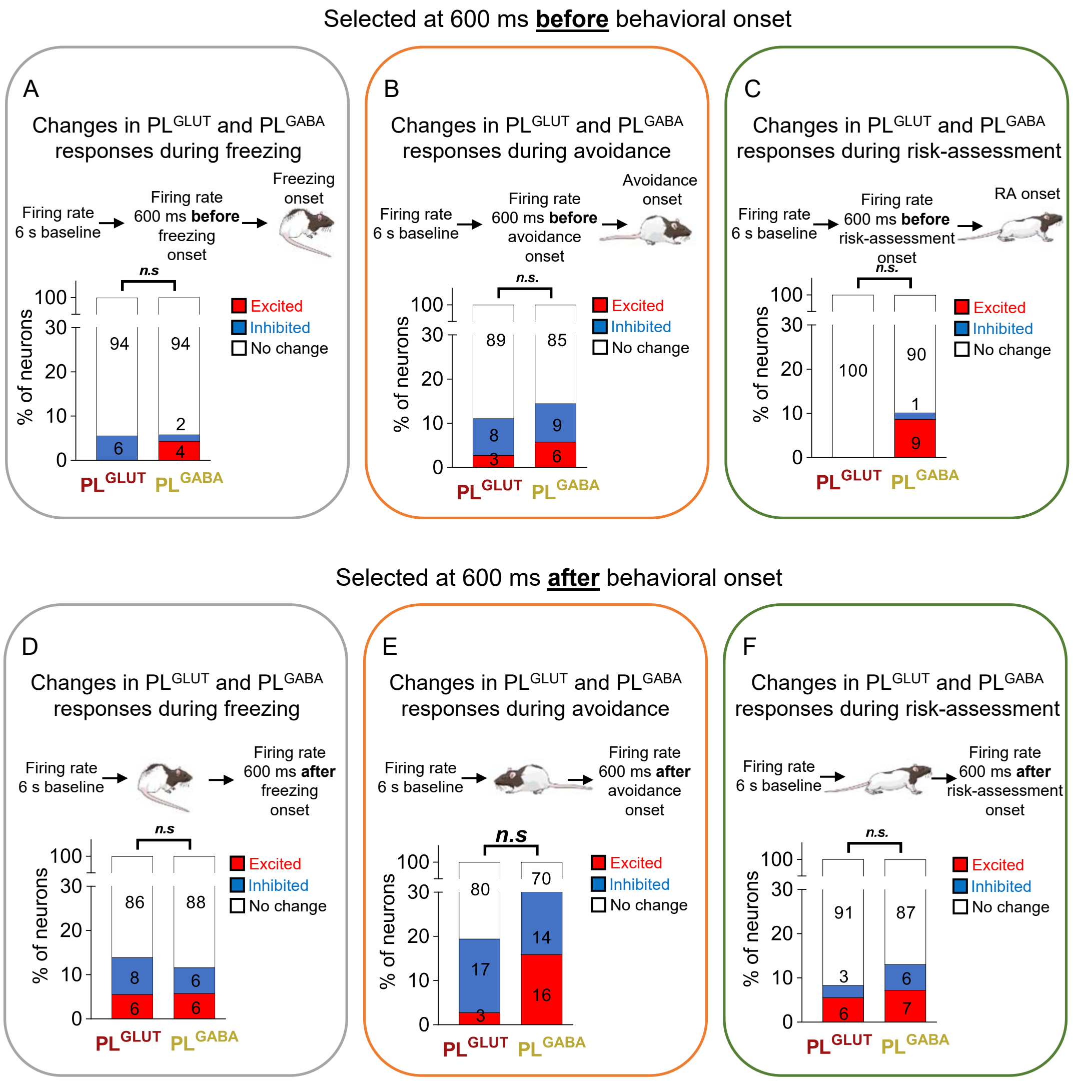

**Supplementary Figure 9. Changes in the firing rate of $PL^{GLUT}$ and $PL^{GABA}$ neurons before or after the onset of freezing, avoidance or risk-assessment behaviors. (A-F)** Both $PL^{GLUT}$ (n = 36) and $PL^{GABA}$ (n = 69) neurons showed the same number and proportion of excitatory and inhibitory responses when aligned at 600 ms before the onset of (A) freezing (Fisher Exact Test, excitatory responses p = 0.549; inhibitory p = 0.270), (B) avoidance (excitatory responses p = 0.6582; inhibitory p >0.999), or (C) risk-assessment (excitatory responses p = 0.919; inhibitory p >0.999) behaviors, as well as when aligned at 600 ms after the onset of (D) freezing (excitatory responses p > 0.999; inhibitory p = 0.688),(E) avoidance (excitatory responses p = 0.0544; inhibitory p = 0.768) or (F) risk-assessment (excitatory responses p = 0.741; inhibitory p = 0.490) behaviors. The threshold used to identify significant differences per neurons was Z-score > 2.58 for excitation and Z-score < -1.96 for inhibition. Stack bar values were compared using Fisher Exact Test, n.s. = non-significant. All statistical analysis details are presented in table S1.

### SUPPLEMENTARY FIGURE 10

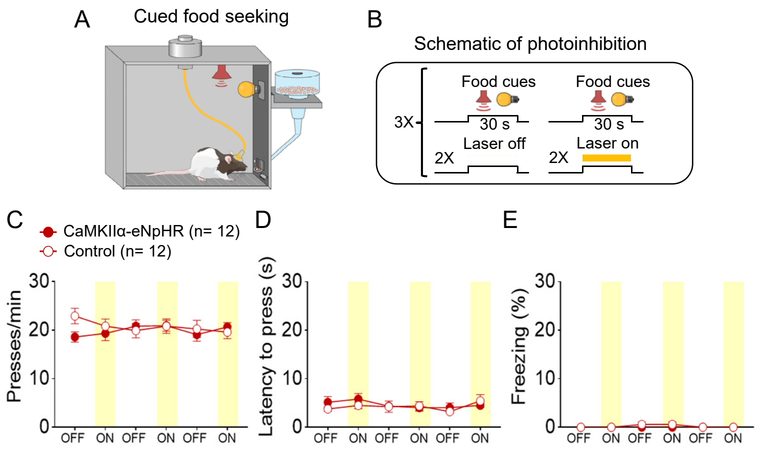

**Supplementary Figure 10. Photoinhibition of $PL^{GLUT}$ neurons did not affect cued food-seeking responses in a neutral context. (A-B)** Schematic and timeline of PL photoinhibition during the cued food-seeking test in a neutral context. **(C-D)** Optogenetic inhibition of $PL^{GLUT}$ neurons (CaMKII-ChR2, dark red circles, n = 12) had no effect on (C) frequency of lever presses ($F_{(5,110)}$ = 1.336, p = 0.254) , (D) latency for the first press ($F_{(5,110)}$ = 0.637, p = 0.671), or (D) freezing responses ($F_{(5,95)}$ = 1.395, p = 0.231), when compared to the control group (eYFP-control virus, white circles, n = 12). Yellow laser illumination (7-10 mW) was delivered for 30 s at cue onset. Data shown as mean ± SEM. Each circle represents the average of two consecutive trials. Two-way repeated-measures ANOVA followed by Bonferroni *post-hoc* test. All statistical analysis details are presented in table S1.

### SUPPLEMENTARY FIGURE 11

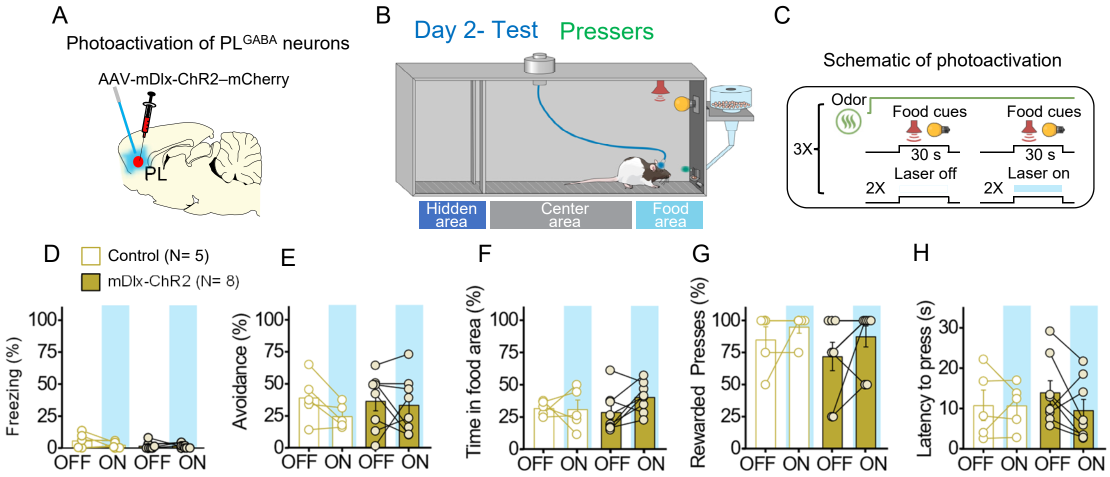

**Supplementary 11. Photoinhibition of $PL^{GABA}$ neurons in *Pressers* does not alter defensive responses and food seeking during conflict (A)** Schematic of AAV-mDlx-ChR2–mCherry virus infusion in PL. **(B-C)** Schematic and timeline of the approach-avoidance conflict test during optogenetic activation of $PL^{GABA}$ neurons. **(D-H)** Photoactivation of $PL^{GABA}$ neurons during the conflict test did not alter rats' behavior (mDlx-ChR2, gold bars, n = 8, Control, white bars, n = 5; Repeated measures ANOVA, Freezing: $F_{(1, 11)} = 2.186$, $p = 0.167$; Avoidance: $F_{(1, 11)} = 0.9568$, $p = 0.349$; Time in food area: $F_{(1, 11)} = 1.798$, $p = 0.207$; Rewarded Presses: Shapiro-Wilk normality test, $p < 0.05$, Wilcoxon test, off vs. on - W = 19, p = 0.312, Mann-Whitney, mDlx-ChR2 vs. control - U=18, p = 0.673; Latency to press: $F_{(1, 11)} = 1.038$, $p = 0.330$). PL neurons were illuminated from cue onset until the animals pressed the lever or from cue onset until the end of the 30 s cues if the animals didn't press the lever ($PL^{GABA}$: 20 Hz; 5 ms pulse width,7-10 mW). Data shown as mean ± SEM. Each bar represents the average of six trials alternated in blocks of 2. All statistical analysis details are presented in table S1.
